# A *studyforrest* extension, MEG recordings while watching the audio-visual movie “Forrest Gump”

**DOI:** 10.1101/2021.06.04.446837

**Authors:** Xingyu Liu, Yuxuan Dai, Hailun Xie, Zonglei Zhen

## Abstract

Naturalistic stimuli, such as movies, are being increasingly used to map brain function because of their high ecological validity. The pioneering *studyforrest* and other naturalistic neuroimaging projects have provided free access to multiple movie-watching functional magnetic resonance imaging (fMRI) datasets to prompt the community for naturalistic experimental paradigms. However, sluggish blood-oxygenation-level-dependent fMRI signals are incapable of resolving neuronal activity with the temporal resolution at which it unfolds. Instead, magnetoencephalography (MEG) measures changes in the magnetic field produced by neuronal activity and is able to capture rich dynamics of the brain at the millisecond level while watching naturalistic movies. Herein, we present the first public prolonged MEG dataset collected from 11 participants while watching the 2 h long audio-visual movie “Forrest Gump”. Minimally preprocessed data was also provided to facilitate the use of the dataset. As a *studyforrest* extension, we envision that this dataset, together with fMRI data from the *studyforrest* project, will serve as a foundation for exploring the neural dynamics of various cognitive functions in real-world contexts.

## Background & Summary

The mechanisms of human brain function in complex dynamic environments is the ultimate mystery that cognitive neuroscience aspires to quest. Most of the existing models on brain function have been obtained from tightly controlled experimental manipulations on carefully designed “artificial” stimuli. However, it remains unclear whether responses to these artificial stimuli can be generalized to ecological scenarios encountered in real-world environments, in terms of quantity, complexity, modality, and dynamics. To address these issues, naturalistic stimuli that encode a wealth of real-life content have become increasingly popular for understanding brain function in ecological contexts. Researchers have achieved significant advances in the areas of human memory, attention, language, emotions, and social cognition using naturalistic stimuli (for recent reviews, please refer to Sonkusare et al., 2019^1^ and Jääskeläinen et al., 2021^2^). Simultaneously, emerging deep learning technologies that could afford multiple levels of representations for naturalistic stimuli are continuously expanding the application of naturalistic stimuli for exploring human brain function^3–6^.

Notably, owing to their dynamics and multimodal content, movies have been successfully utilized as naturalistic stimuli to examine the mechanism by which the brain processes diverse psychological constructs and dynamic interactions. Functional magnetic resonance imaging (fMRI) is commonly employed to measure brain activity while watching a movie. In particular, the pioneering *studyforrest* and other naturalistic neuroimaging projects have released multiple fMRI datasets, collected from participants who had watched movie clips^7–11^. However, fMRI measures the relatively sluggish blood-oxygenation-level-dependent signal, therefore falling short of characterizing the complex neural dynamics underlying the cognitive processing of dynamic movies. In contrast, magnetoencephalography (MEG) measures the magnetic fields generated by neuronal activity on a millisecond time scale. Thus, it has great potential to pry open neural dynamics in processing naturalistic stimuli. Several studies have leveraged MEG to investigate brain activity for naturalistic movie stimuli in a short period (≤ 20 min)^12–16^. However, there is still a dearth of publicly accessible MEG recordings for naturalistic stimuli, especially prolonged MEG recordings for dynamic movies that are more likely to capture the temporal dynamics of regular functional brain states that occur in everyday life, and further contribute to unraveling human brain function in ecological contexts.

Herein, we present an MEG dataset obtained while watching the 2 h long audio-visual movie “Forrest Gump” (R. Zemeckis, Paramount Pictures, 1994). The recordings measure brain activation with a temporal resolution at the millisecond level, thus providing a timely and efficient extension to the *studyforrest* dataset. Specifically, MEG data were collected from 11 participants while they were watching the Chinese-dubbed movie “Forrest Gump” in eight consecutive runs, each lasting for roughly 15 min. High-resolution structural MRI was additionally acquired for all participants, thereby allowing the incorporation of the detailed anatomy of the brain and head in the source localization of MEG signals. Together with the raw data, preprocessed MEG and MRI data with standard pipelines were also provided to facilitate the use of the data. Considering MEG and fMRI are complementary to each other, synergy between our present MEG recordings and fMRI data from the *studyforrest* project will provide a valuable resource to study brain function in the real-life contexts. We believe the dataset is suitable for addressing many questions pertaining to the neural dynamics of various aspects, including perception, memory, language, and social cognition.

## Methods

### Participants

A total of 11 participants (mean±SD age: 22±1.7 years, 6 female participants) from the Beijing Normal University, Beijing, China, volunteered for this study. They completed both the MEG and MRI sessions. All participants were right-handed, native Chinese speakers, with normal hearing and normal or corrected-to-normal vision. None of them had ever watched the film “Forrest Gump” before, except one who had watched some clips, however not the entire movie. Of the 10 participants, four had heard about the movie plot, while others did not. The study was approved by the Institutional Review Board of the Faculty of Psychology, Beijing Normal University. Written informed consent was obtained from all participants, prior to their participation. All participants provided additional consent for sharing their anonymized data for research purposes.

### Procedures

Fig. 1 depicts the overall flow of data collection and preprocessing. Prior to data acquisition, all participants completed a questionnaire on their demographic information and familiarity with the movie “Forrest Gump”. The data acquisition consisted of two sessions for each participant, namely, one MEG session to record their neural activities during movie watching and an MRI session with a T1-weighted (T1w) scan to measure the brain structure for the spatial localization of the MEG signal. The MRI scan immediately followed the MEG session for all participants, except for sub-07 and sub-11, who finished their MRI session a week later.

**Fig. 1.**
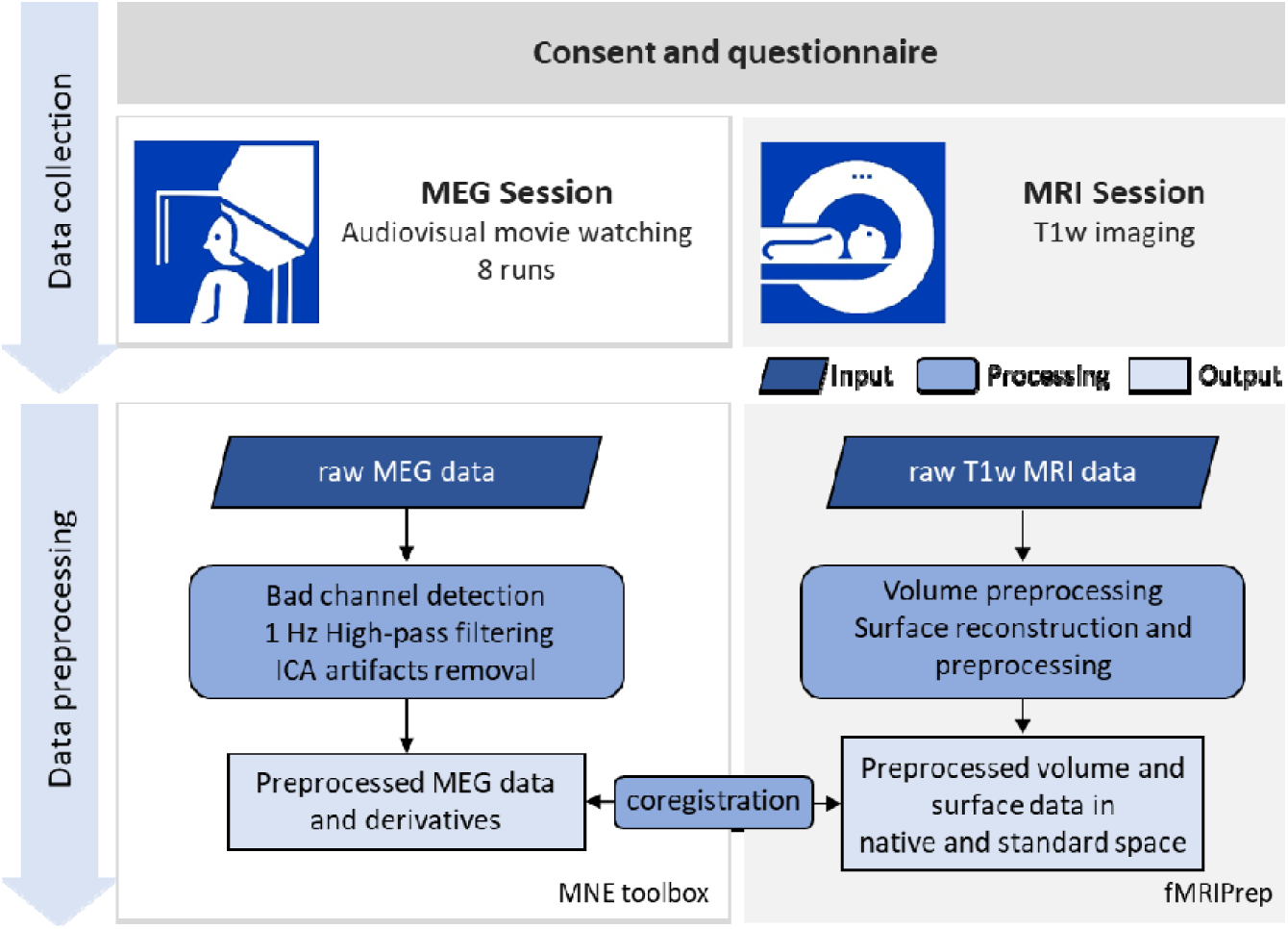
Schematic of the data collection and preprocessing procedure. Data collection comprised of one MEG session followed by one MRI session. The neuromagnetic signals were recorded with a whole-scalp-covering MEG while the participants watched the audio-visual movie “Forrest Gump”. An anatomical T1w imaging was acquired in the MRI session. The raw MEG data and MRI data were preprocessed with MNE and fMRIPrep toolbox, respectively. The MEG-MRI coregistration was performed on the preprocessed data.

### Stimulus material and presentation

The audio-visual stimuli were generated from the Chinese-dubbed “Forrest Gump” DVD, released in 2013 (ISBN: 978-7-7991-3934-0). The movie was split into eight segments, each of which lasted for approximately 15□min. The stimuli were initially obtained by concatenating all original VOB files from the DVD release into one MPEG-4 file, using FFmpeg (https://ffmpeg.org). The concatenated MPEG-4 file contained a video stream and a Chinese-dubbed audio stream, which was down-mixed from multi-channel to 2Channel stereo. The stimuli were then divided into eight segments using Adobe Premier software (Adobe Premiere Pro CC 2017, Adobe, Inc., San Jose, CA, USA). Each segment conformed to the following specifications: video codec=avc1, display aspect ratio=4:3, resolution=1024×768 pixels, frame rate=25 FPS, color space=YUV, video bit depth=8 bits, audio codec=mp4a-40-2, audio sampling rate=48.0 kHz, and audio channels=2. Each successive segment began with a 4 s repetition of the end of the previous segment. It should be noted that the Chinese-dubbed “Forrest Gump” is an abridged edition of the German version; some segments in the German version are not included in the Chinese version. To align with the stimuli of the *studyforrest* dataset as much as possible, a short clip from the German-dubbed DVD released in 2011 (EAN: 4010884250916) was added to our stimuli. Table 1 summarizes the sources of the MEG stimuli from both the Chinese and German versions. Fig. 2 characterizes the alignment between the MEG stimuli (current study) and fMRI stimuli (*studyforrest* project) for each run.

**Table 1.**
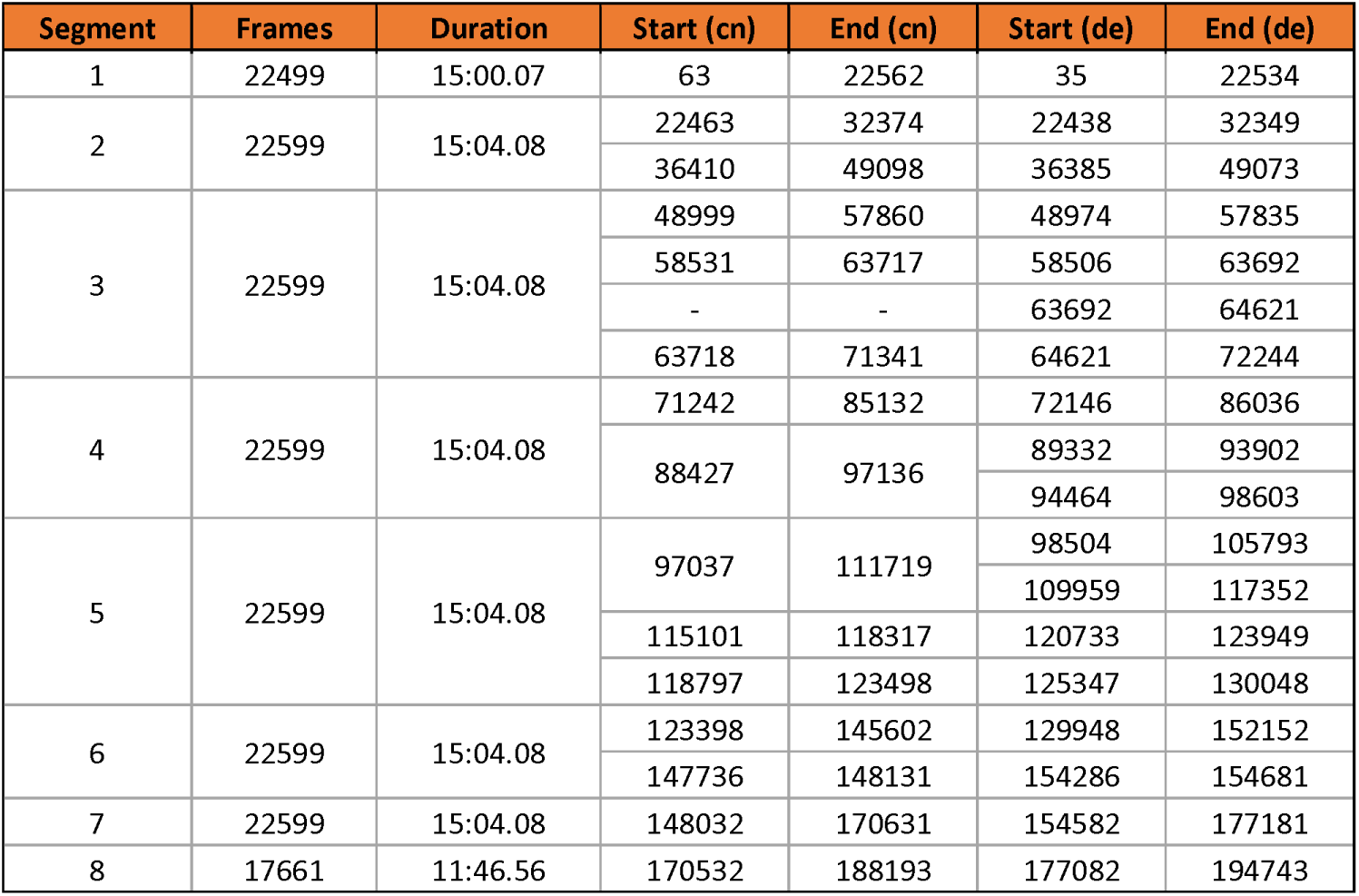
Stimulus sources from the Chinese (cn) and German (de) version of “Forrest Gump”. The start time and end time are different for the same movie clip in the two versions. Additionally, to better align with the existing *studyforrest* dataset, a clip from the German version (frames from 63692 to 64621, ∼37 sec) was added into segment 3 after removing the vocal sound track of the clip. Specifically, the visual stream remains identical to the German version while the audio stream only contains the background sound track.

| Segment | Frames | Duration | Start (cn) | End (cn) | Start (de) | End (de) |
| --- | --- | --- | --- | --- | --- | --- |
| 1 | 22499 | 15:00.07 | 63 | 22562 | 35 | 22534 |
| 2 | 22599 | 15:04.08 | 22463 | 32374 | 22438 | 32349 |
|  |  |  | 36410 | 49098 | 36385 | 49073 |
| 3 | 22599 | 15:04.08 | 48999 | 57860 | 48974 | 57835 |
|  |  |  | 58531 | 63717 | 58506 | 63692 |
|  |  |  | - | - | 63692 | 64621 |
|  |  |  | 63718 | 71341 | 64621 | 72244 |
| 4 | 22599 | 15:04.08 | 71242 | 85132 | 72146 | 86036 |
|  |  |  | 88427 | 97136 | 89332 | 93902 |
|  |  |  |  |  | 94464 | 98603 |
| 5 | 22599 | 15:04.08 | 97037 | 111719 | 98504 | 105793 |
|  |  |  |  |  | 109959 | 117352 |
|  |  |  | 115101 | 118317 | 120733 | 123949 |
|  |  |  | 118797 | 123498 | 125347 | 130048 |
| 6 | 22599 | 15:04.08 | 123398 | 145602 | 129948 | 152152 |
|  |  |  | 147736 | 148131 | 154286 | 154681 |
| 7 | 22599 | 15:04.08 | 148032 | 170631 | 154582 | 177181 |
| 8 | 17661 | 11:46.56 | 170532 | 188193 | 177082 | 194743 |

**Fig. 2.**
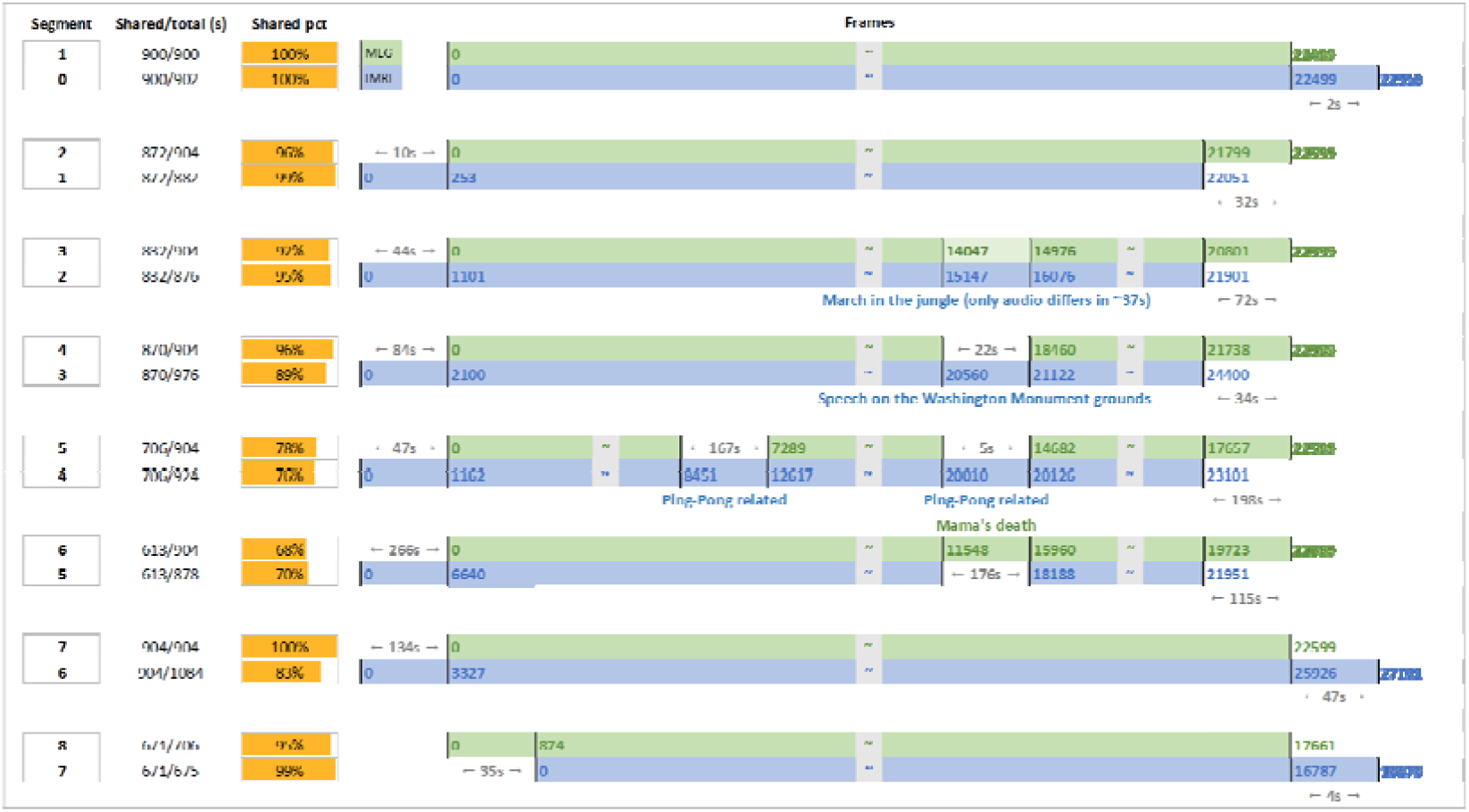
Stimulus alignment between the MEG and fMRI dataset. Both the matched and the mismatched parts between the MEG (green) and fMRI (blue) stimuli were marked in the timeline for each segment (i.e., run). The mismatched segments were particularly annotated with frame number, duration, and a brief scene description. The shared fraction was shown in both time length and percentage (the yellow bar). For display purpose, the length of segments are not represented in the real scale.

The detailed configuration of the MEG system is illustrated in Fig. 3. The movie stimuli were presented using Psychophysics Toolbox Version 3^17^ in MATLAB 2016 (MathWorks, Natick, MA, USA), which drives the PC hardware directly to generated stimuli. The audio stimuli were generated with a standard PCI sound card (Sound Blaster Audigy 5/Rx) and then amplified with an insert earphone. The insert earphone comprised of a small driver unit (E-A-R-TON transducer; 3M, Minnesota, USA), a thin plastic tube (Tygon B-44-4x tubing; Saint Gobain Tygon division), and an earplug (ER3-14A; Etymotic Research, Inc., Elk Grove Village, IL, United States). The visual stimuli were generated with a standard PCI graphics card (GeForce GT620; NVIDIA, Santa Clara, California, USA) and projected onto a screen in full-screen mode via a DLP projector (NP63+; NEC, Tokyo, Japan) with 1024×768-pixel resolution. The participants watched the visual stimuli on a rear projection screen through mirror reflection (visual field angles=31.17°×23.69°; viewing distance=751 mm). The participants were instructed to watch the movie, without performing any other tasks and to keep still as best as possible.

**Fig. 3.**
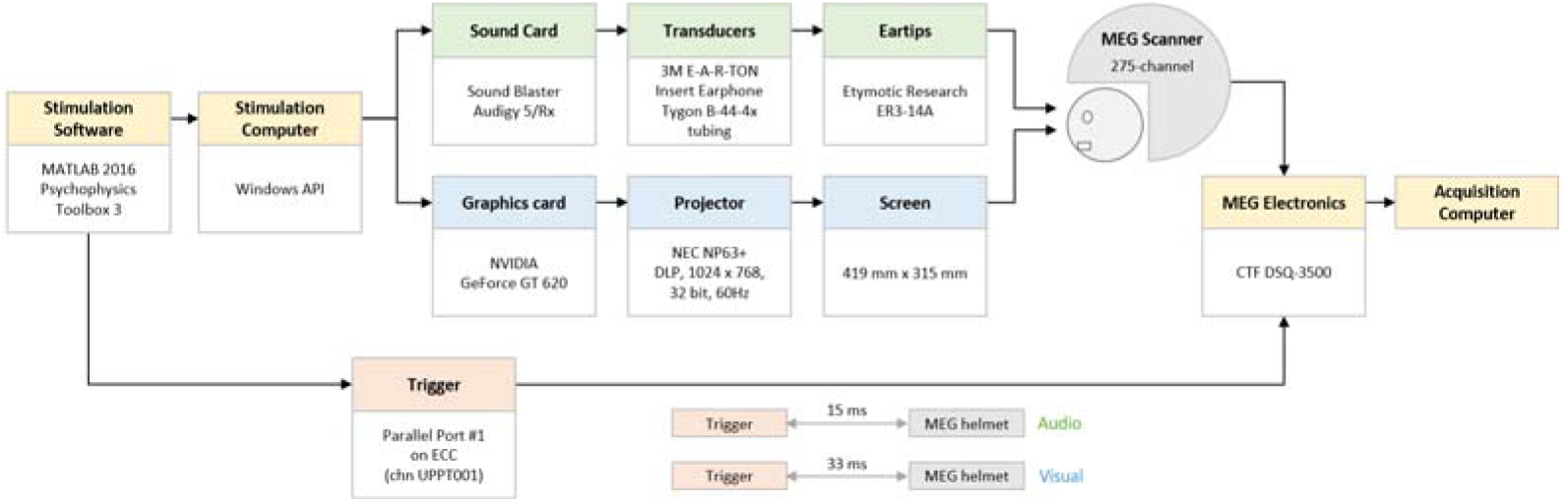
Configuration diagram of the MEG system. Both the audio and visual stimulus systems were custom made as a whole, though most of their components are commercially available. The average latencies between the time at which the stimulation was received and the time of trigger pulse were 33 ms and 15 ms for the visual and audio stimuli, respectively.

The average latencies between the stimulus initiation and the actual receipt of the stimulus was 33 ms and 15 ms for the visual and audio stimuli, respectively. The latencies were measured independently from the MEG movie experiment. A standard pulse stimulus (visual: white patch, audio: pure tone) was requested to be presented by Psychophysics Toolbox, while the corresponding onset trigger was sent to the trigger channel. A photo (visual) / acoustic (audio) sensor connected to one of the MEG Analog-to-Digital Converter (ADC) channels, placed near the MEG helmet, was used to detect the stimuli. The latencies were then calculated as the difference between the recorded time of the ADC channel and the trigger channel. The latencies were measured multiple times and the average latencies were calculated for visual and audio stimulation respectively.

### MEG data acquisition

MEG data were recorded using a 275-channel whole-head axial gradiometer DSQ-3500 MEG system (CTF MEG, Canada) at the Institute of Biophysics, Chinese Academy of Sciences, Beijing, China. Three channels (i.e., MLF55, MRT23 and MRT16) were out of service due to failure of sensors. The neuromagnetic signals were recorded in continuous mode at a sampling rate of 600 Hz, without online digital band filters. A third-order synthetic gradiometer was employed to remove far-field noise. The precise timing of each frame was recorded. After each frame was requested to be presented to the participants, we recorded a trigger pulse lasting for five samples in the stimulus channel UPPT001. The beginning of the movie was indicated with a value of 255. Owing to the limited bit-width of the stimulus channel, the frame number could not be marked with an accurate value > 20,000. The frame numbers were therefore marked as the ceiling of the timestamp of that frame divided by 10, resulting in step-like increasing marker values starting from 1 with a step width of 250 (25 fps * 10 s).

At the beginning of each session, three HPI coils were attached to the participants’ nasion (NAS), left preauricular (LPA), and right preauricular (RPA) points to continuously measure their head position in the MEG helmet. A customized wooden chin-rest supporter was introduced to prevent possible head movements. The MEG session consisted of eight runs, with each run playing one movie segment. Eight segments were played chronologically. The participants took a self-paced break between runs. Following the completion of the MEG scan, the participants underwent an anatomical T1w scan. The HPI coils were replaced with three customized MRI-compatible vitamin E capsules in the MRI scan to provide spatial reference for the spatial alignment between the MEG and MRI data.

### MEG data preprocessing

MEG data processing was performed offline using the MNE-Python v0.22 package^18^. The MEG preprocessing pipeline was conducted at the run level (Fig. 1). First, the bad channels were detected and marked. As a result, no bad channels were identified in all acquisitions except two in run-05 of sub-05. Second, a high-pass filter of 1 Hz was applied to remove possible slow drifts from the continuous MEG data (Fig. 4a). Finally, artifact removal was performed using an independent component analysis (ICA) with FastICA wrapped in the MNE. Artifact-ICs were classified mainly using the spatial topography and time course information following the guide provided by Bishop & Busch (2015)^19^. The number of independent components (IC) was set to 20. Two raters (i.e., X.L. and Y.D.) manually identified the head movement, eye movement, eye blinks, and cardiac artifacts (Fig. 4b). On average, 3.21 ICs (SD: 0.85) were classified as artifacts. The denoised MEG data were eventually reconstructed from all the non-artifact components and residual components (Fig. 4c). Both the raw and preprocessed data were provided in the released dataset.

**Fig. 4.**
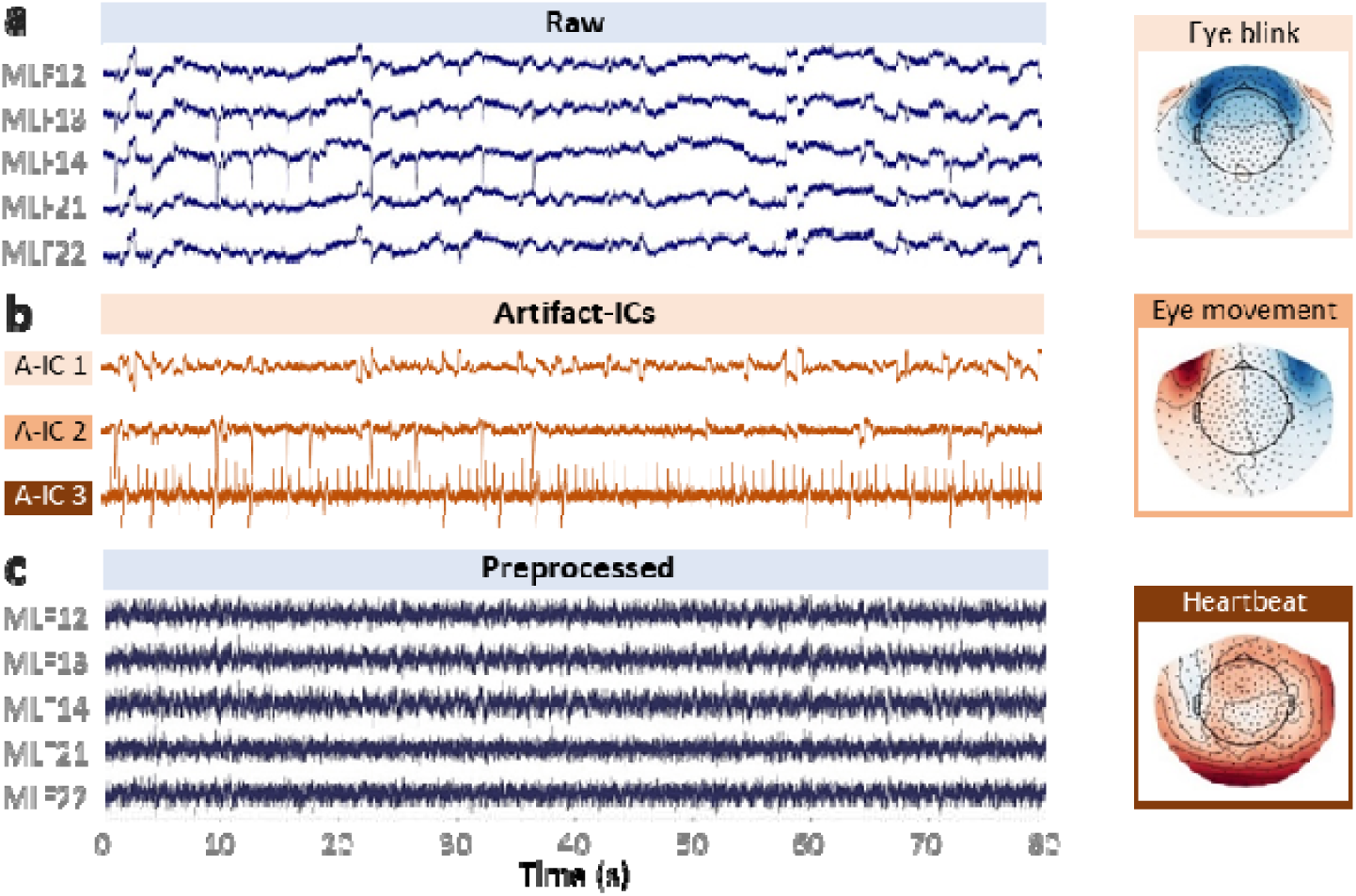
Typical artifact-ICs and MEG signals from the pre- and post-preprocessing data. **(a)** MEG signals of example channels from the raw, i.e., pre-preprocessing data. **(b)** Timeseries and scalp field distribution of three typical artifact-ICs (A-ICs), namely A-IC 1 for eye blink, A-IC 2 for horizontal eye movement, and A-IC 3 for heartbeat. **(c)** MEG signals of example channels from the preprocessed data. Data from the run-04 of sub-04 were used for this illustration.

### MRI data acquisition and preprocessing

High-resolution anatomical MRI was collected for each participant using a 3T SIEMENS Prisma^fit^ scanner (Siemens Healthcare GmbH, Erlangen, Germany), with a 20-channel headneck coil. All participants underwent a T1w scan with a 3-D magnetization-prepared rapid gradient-echo pulse sequence with identical parameters (TR=2530 ms, TE=1.26 ms, TI=1100 ms, flip-angle=7°, 176 sagittal slices, slice thickness=1 mm, matrix size=256×256, and voxel size=1.0□×1.0 mm), except that sub-01 was scanned with slightly different parameters (TR=2200 ms, TE=3.37 ms, TI=1100 ms, flip-angle=7°, 192 sagittal slices, slice thickness=1 mm, matrix size=224×256, and voxel size=1.0□×1.0 mm). Earplugs were used to attenuate the scanner noise. A foam pillow and extendable padded head clamps were utilized to restrain the head motion.

The raw DICOM files of T1w images were converted to NIFTI files using dcm2niix (https://github.com/rordenlab/dcm2niix). The T1w images were then minimally preprocessed using the anatomical preprocessing pipeline from fMRIPrep v20.2.1, with default settings^20^. In brief, the T1w data were skull-stripped and corrected for intensity nonuniformity with ANTs and N4ITK^21^. Brain surfaces were reconstructed using FreeSurfer^22^. Spatial normalization to both MNI152NLin6Asym and MNI152NLin2009cAsym was performed through nonlinear registration with ANTs, using the brain-extracted versions of both T1w volume and template.

### MEG-MRI coregistration procedure

To reconstruct the source of MEG sensor signals, MEG data were co-registered with the high-resolution anatomical T1w MRI data for each participant. The NAS, LPA, and RPA points marked in both MEG and MRI sessions were used as fiducial points for the alignment of the MEG and MRI data. Specifically, following the generation of a high-resolution head surface using MNE make_scalp_surfaces based on FreeSurfer reconstruction, we performed MEG-MRI coregistration for each participant in the MNE COREG GUI^18^. First, the three fiducial points were manually pinned on the MRI-reconstructed head surface. An iterative algorithm (nearest-neighbor calculations) was then run to align the MEG and MRI coordinates. The co-registration was refined by manual adjustment. The results showed that the average distances between the three fiducials in the coregistered MEG and MRI coordinate systems were 0.96 mm, 4.22 mm, and 4.90 mm for NAS, LPA, and RPA, respectively. Both the MRI-fiducials files and MEG-MRI coordinate transformation files were included in the released data.

### Data Records

The dataset can be accessed at OpenNeuro (dataset accession number: ds003633, version 1.0.3, https://openneuro.org/datasets/ds003633/versions/1.0.3)^23^. The facial information was removed from the published dataset using pydeface (https://github.com/poldracklab/pydeface) to ensure anonymity. The data was organized according to the MEG-Brain Imaging Data Structure^24^ v1.4.0 using the MNE-BIDS v0.8 toolbox^25^ (Fig. 5). Besides dataset and participant description files, the data were sorted into different directories, including “sub-<participant_id>,” “derivatives,” and “code” directory for raw data from each participant, preprocessed data, and the code used for stimuli presentation, MEG and MRI data preprocessing, respectively (Fig. 5a).

**Fig. 5.**
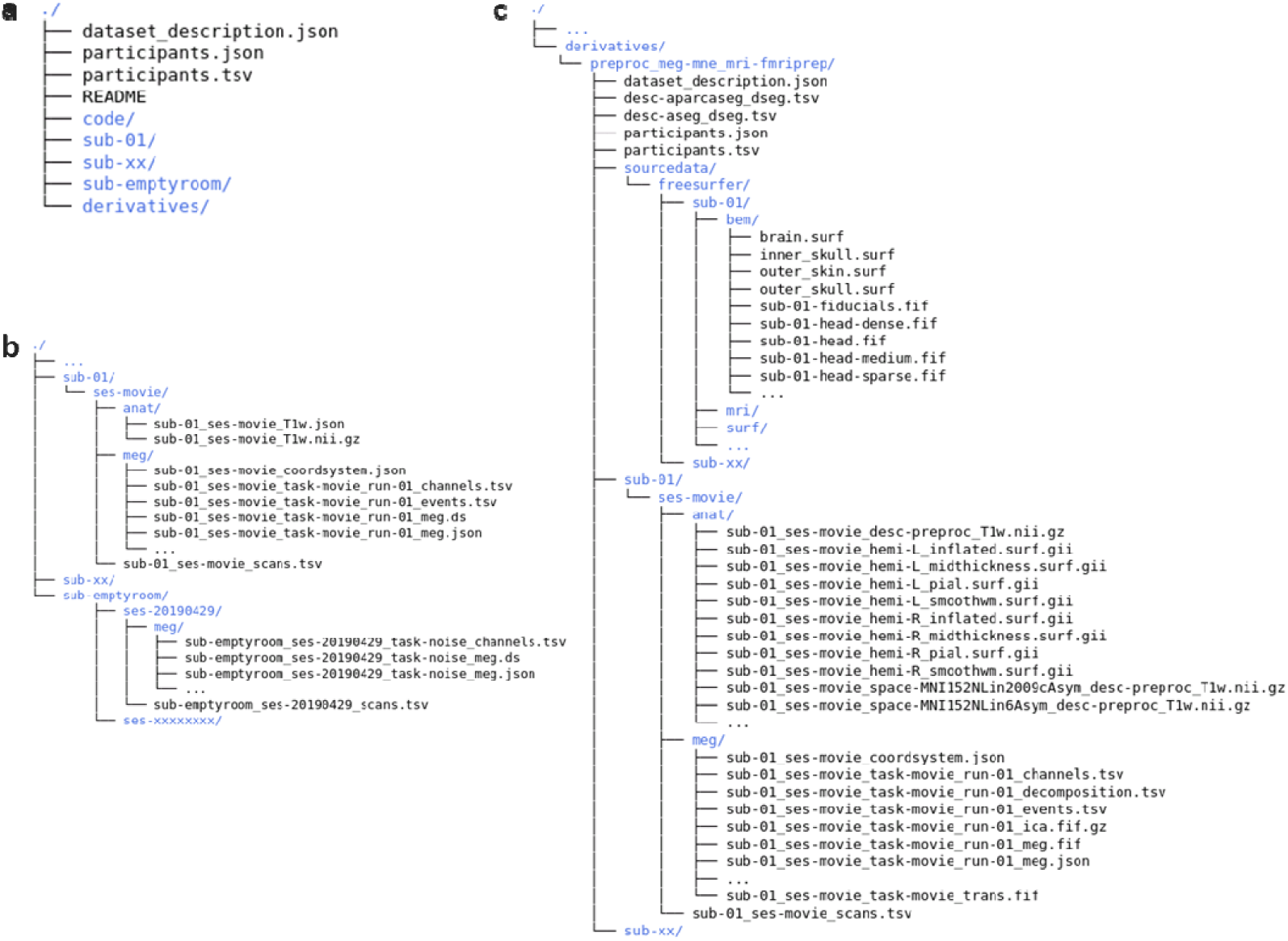
File structure of the dataset. **(a)** File structure of the project directory. **(b)** File structure of the raw data for each individual participant. **(c)** File organization of the derived (preprocessed) data.

### Raw data

The raw data of each participant were stored separately in the “sub-<participant_id>“ folders (Fig. 5b), consisting of two subfolders, namely “anat” and “meg”. The T1w MRI data (“*T1w.nii.gz”) and associated sidecar json files were located in the “anat” folders. The raw MEG data were provided as CTF ds files (“*_meg.ds”) for each run, and located in the “meg” folder along with sidecar json files. In addition, “*_channel.tsv” files with MEG channel information, “*_events.tsv” files with the presentation timing of stimuli frames, and “*coordsystem.json” files with coordinate system information of the MEG sensors were included in the “meg” folder.

In parallel with the “sub-<participant_id>“ directories, a “sub-emptyroom” directory hosted empty-room MEG measurements, which recorded the environmental noise of the MEG system. The empty-room measurements lasted for 34 s and were acquired on each data acquisition day, except for the day 20190603.

### Preprocessed data

All preprocessed data were deposited in the “preproc_meg-mne_mri-fmriprep” subdirectory under the “derivatives” (Fig. 5c). The preprocessed data of each participant were separately saved in the “sub-<participant_id>/ses-movie” directory, which contains two subfolders, namely “anat” and “meg”. The “anat” folder comprised the preprocessed MRI volume, reconstructed surface, and other associations, including transformation files. The “meg” folder included preprocessed MEG recordings, including “*_meg.fif.gz”, “*_ica.fif.gz” and “*_decomposition.tsv”, and “*_trans.fif” for the preprocessed data, ICA decomposition, and MEG-MRI coordinate transformation, respectively. In addition, the FreeSurfer surface data, the high-resolution head surface (“freesurfer/sub-<participant_id>/bem/*”), and the MRI-fiducials (“freesurfer/sub-<participant_id>/bem/*fiducials.fif”) were provided in “freesurfer/sourcedata” directory for MEG-MRI coregistration.

### Technical Validation

We assessed the data quality of both the raw and preprocessed data using four measures as follows: head motion magnitude, stimuli-induced time-frequency characteristics, homotopic functional connectivity (FC), and inter-subject correlation (ISC).

### Motion magnitude distribution

Head movements during MEG scans are one of the significant factors that degrade both sensor- and source-level analyses. Herein, we calculated the motion magnitude for each sample as the Euclidian distance between the current and the initial head position while the movie segment began playing. The head motion across all runs and all participants were summarized for each of the three fiducials (NAS, LPA, and RPA) to provide an overview of the head movement of the dataset. As shown in Fig. 6, motion magnitude of 95% of the samples had head motions lower than 3.43 mm, 4.11 mm, and 3.87 mm for NAS, LPA, and RPA, respectively.

**Fig. 6.**
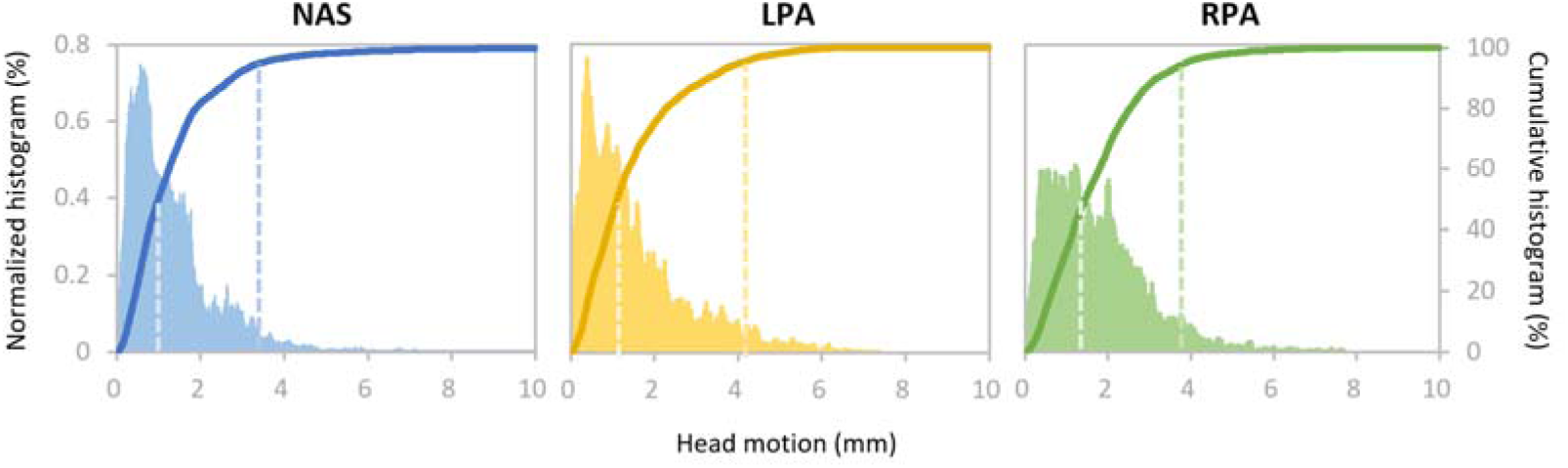
Ensemble distribution of head motion magnitude across all runs and all participants. The density and accumulative histogram of motion magnitude of all samples from all acquisitions for three fiducials (NAS, LPA, and RPA) have been plotted. The dashed lines indicate 50% and 95% of the cumulative density. Left Y-axis: normalized histogram; Right Y-axis: cumulative histogram.

Furthermore, 50% of the samples had head motions smaller than 0.99 mm, 1.10 mm, and 1.46 mm for NAS, LPA, and RPA, respectively. These findings indicated low head motion magnitude on average. The head motion magnitudes of each participant are presented in Supplementary Fig. 1.

### Time-frequency characterization of brain activity

Herein, we validated whether MEG recordings could successfully detect the change in stimuli-induced brain activity. Because the movie stimuli do not have explicit condition structures as in conventional design, we selected two exemplar movie clips, within which the audio or visual features showed pronounced changes to examine if the expected change in MEG signals could be detected at the related sensors. In one clip (Seg 3: frame 15864±125 [10:35±5 sec], the scene where Gump was marching during the Vietnam War), the audio features changed significantly (a vocal voice developed from background music), whereas the visual features were stable. In contrast, the other clip (Seg 1: frame 21768±125 [14:30±5 sec], the scene where young Gump and Jenny were sitting on the tree waiting for stars), comprised of stable audio features, whereas the visual features changed from landscape to human figures. Validation was performed according to the following procedure^26–28^: first, time-frequency analysis with Morlet wavelets was conducted for each sensor in the occipital and temporal lobes. The baseline was set to 1000 ms before the change points of the audio or visual features, and the baseline mean was subtracted for each channel. Second, the time-frequency representations were averaged across the participants. Finally, the time-frequency representations were averaged across the sensors from the occipital and temporal lobes. As shown in Fig. 7, the time-frequency representations from the occipital sensors, and not the temporal sensors, were locked with changes in visual features. The opposite pattern was observed for the audio feature changes in the stimuli. The results demonstrated that current MEG data could accurately detect stimulus-induced brain activity.

**Fig. 7.**
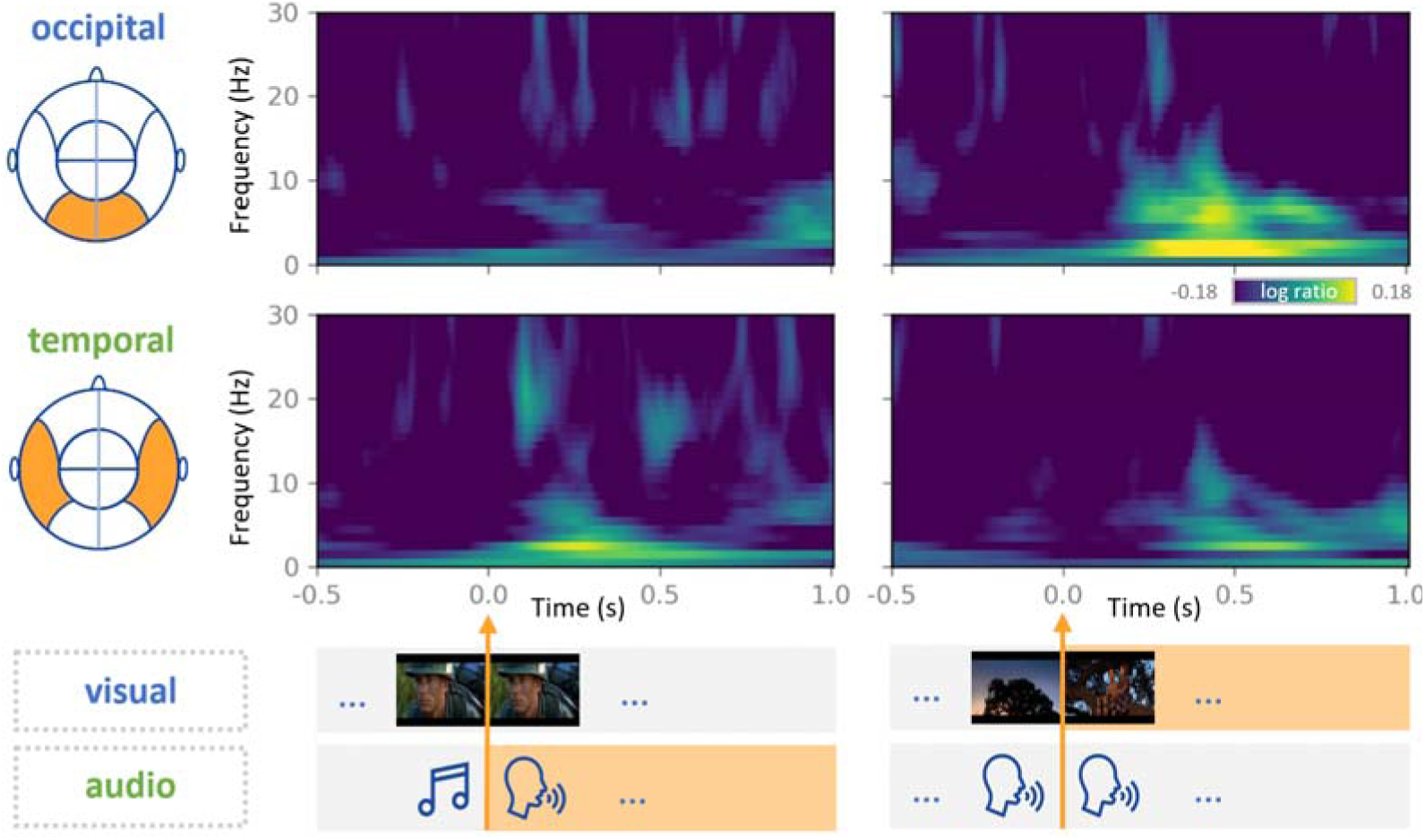
Time-frequency characterization of the brain activity for stimuli. MEG signals from two movie clips with pronounced changes in either audio or visual features of the stimuli have been examined. The time-frequency (TF) spectrum is shown for each condition with a unit of log ratio between the TF spectrum calculated from the signal of interest and that from the baseline signal (1000 ms before the onset). The occipital sensors (top) display significant signal changes with a change in the visual features of the stimuli. The temporal sensors (bottom) display significant changes with a change in the audio features.

### Homotopic functional connectivity

A basic principle of the brain’s functional architecture is that FC between inter-hemispheric homologs (i.e., homotopic regions) is particularly stronger than other interhemispheric (i.e., heterotopic) FCs^29,30^. Herein, we tested if MEG data for dynamic movies at the sensor level could reveal strong homotopic FC. First, the absolute envelope amplitude of the MEG signal for each sensor (i.e., channel) was calculated using the Hilbert transform and then down-sampled to 1 Hz. Second, the homotopic FC between a certain sensor and its homotopic sensor was calculated as the Pearson correlation coefficient. For comparison, the heterotopic FC was also calculated for each sensor as the average correlation between it and all its heterotopic sensors. Finally, the homotopic and heterotopic FC values were averaged across all runs and all participants. The homotopic FCs were generally stronger than the average heterotopic FCs (Fig. 8a). Importantly, the high homotopic FC primarily appeared at the sensors located in the occipital and temporal cortices, thereby indicating strong couplings driven by the movie stimuli (Fig. 8b).

**Fig. 8.**
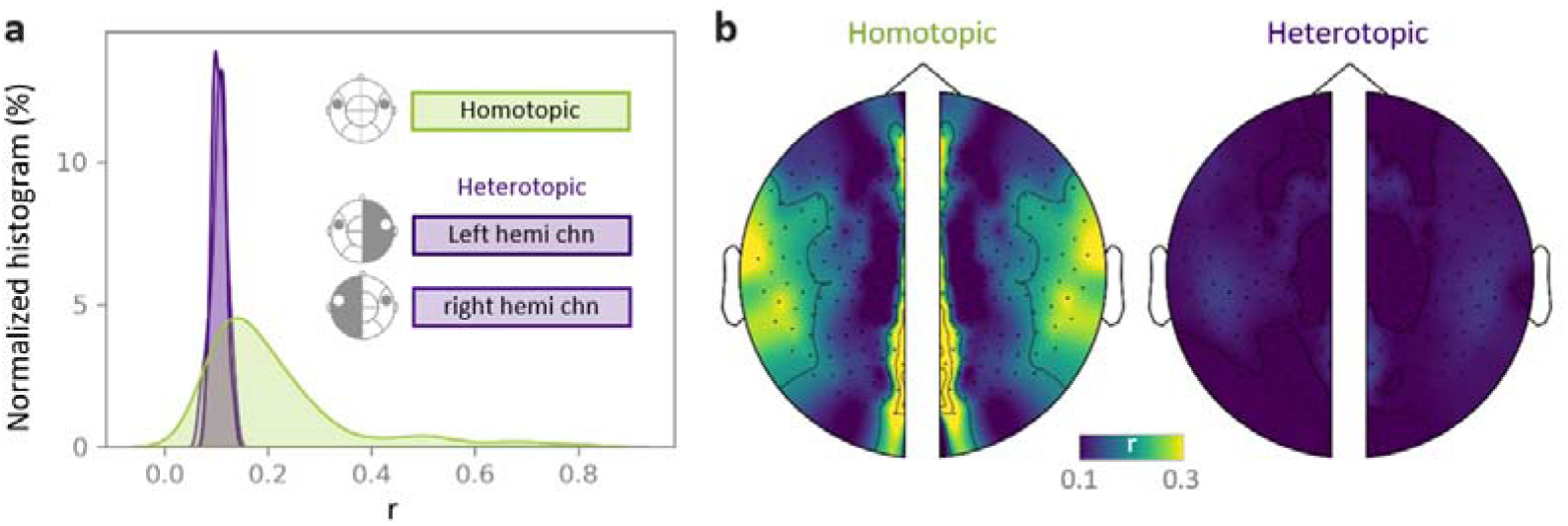
Sensor-level homotopic functional connectivity (FC) is stronger than heterotopic FC. **(a)** Density histogram of the sensor-level homotopic FC and heterotopic FC pooled across all runs and participants. **(b)** Topographic maps of the sensor-level homotopic (left) and heterotopic (right) FCs averaged across all acquisitions. Identical homotopic FC values are displayed for the corresponding homotopic sensors from the two hemispheres. Sensors in the central axis with no corresponding homotopic sensors were not included in the analysis.

### Inter-subject correlation

ISC analysis uses the brain responses of a subject to naturalistic stimuli as a model to predict the brain responses of other subjects^31^. Numerous studies have demonstrated that visual and auditory cortices display significant ISC while watching audio-visual movies. We validated if a high ISC could be detected in our MEG data. For simplicity, the ISC analysis was conducted at the sensor-level. The MEG recordings captured neural oscillations at different frequency bands^32^. Therefore, the ISC was calculated in five bands (delta: 1-4 Hz, theta: 4-8 Hz, alpha: 8-13 Hz, beta: 13-30 Hz, and gamma: 30-100 Hz). First, the MEG signal was filtered for each band.

Second, the absolute envelope amplitude of each band was calculated via the Hilbert transform and down-sampled to 1 Hz. Third, for each frequency band, a leave-one-participant-out ISC was calculated for the left participant as the temporal correlation between the envelope amplitude from the participant and the average of other participants. Finally, the mean ISC was calculated by averaging the ISC across all participants. As shown in Fig. 9, sensors with higher ISC were located near the visual and audio cortices in the preprocessed data, which reportedly displayed high ISC during movie watching in previous studies^14,33–35^. In particular, the high ISC predominantly occurred in the delta, theta, and alpha bands, consistent with previous studies^14,34,35^. Together, the dataset demonstrated good validity in detecting ISC.

**Fig. 9.**
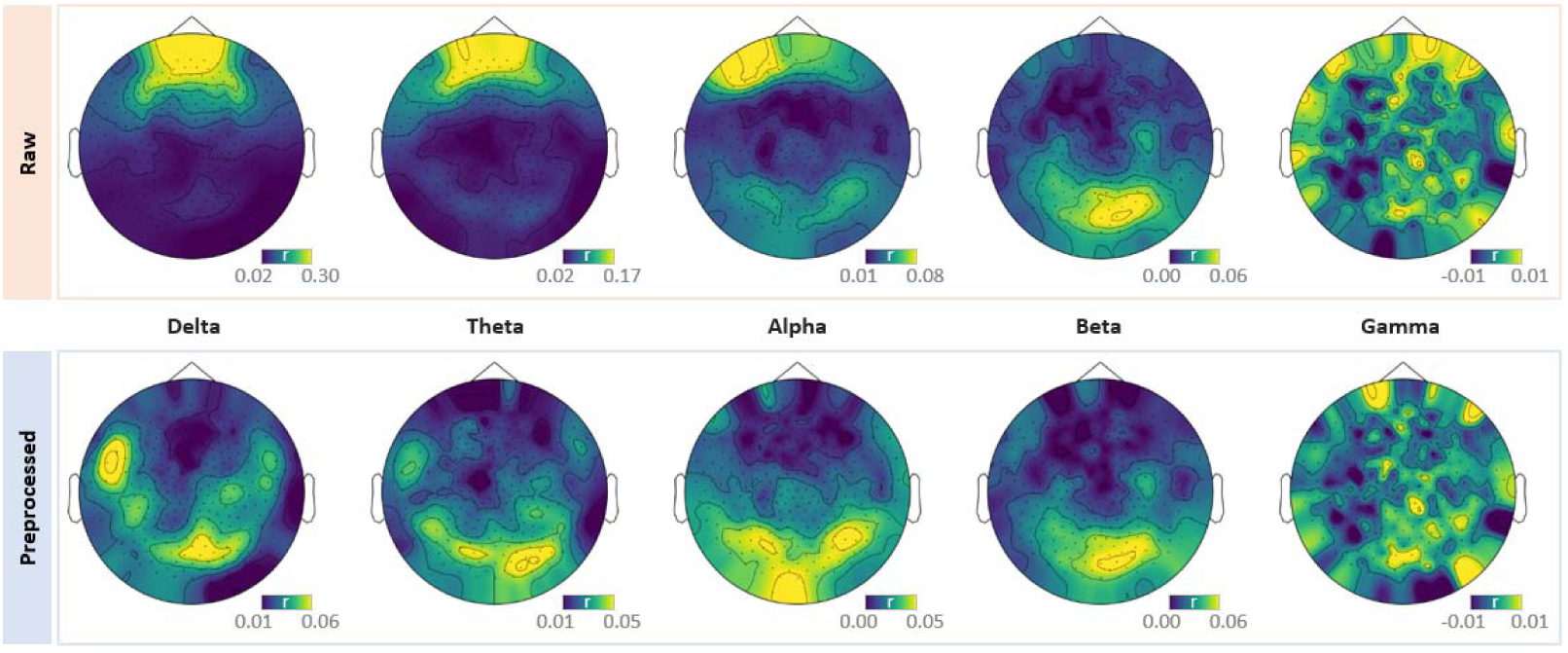
Topographic maps of ISC in different frequency bands derived from both the raw and preprocessed MEG data. A high ISC occurs near the visual and audio cortices in the delta, theta and alpha bands in the preprocessed data. The high ISC around the orbital area is only observed in the raw data but not in the ICA denoised data.

In addition, the high ISC around the orbital area in the raw data implies a synchronized eye movement across participants, which may have been generated by the participants’ high engagement with the movie^36^. These similar eye movement patterns produced similar electrooculography artifacts, which in turn caused high ISC around the orbital area in the raw data. The fact that the high ISC around the orbital area is only observed in the raw data but not in the ICA denoised data testified this speculation. Therefore, these results also reflect a good validity of our preprocessing protocol.

### Usage Notes

We presented the first public MEG dataset for a full-length movie. The MEG signals were recorded while the participants watched the 2 h long Chinese-dubbed audio-visual movie “Forrest Gump”. The dataset provided a versatile resource for studying information processing in real-life contexts. First, MEG data could be independently used to study the neural dynamics of sensory processing and higher-level cognitive functions under real-life conditions. Second, as a *studyforrest* extension, the dataset could be integrated with publicly available fMRI data from the *studyforrest* project. The fusion of fMRI and MEG may shed new light on the relationship between spatially localized networks observed in fMRI and the MEG-derived temporal dynamics. Moreover, our massive MEG recordings enable the direct training of deep neural networks (DNNs) with neural activity patterns. In contrast to the DNNs that were usually trained with stimuli without referring to any neural representation, this kind of brain-constrained DNNs would act more like the human brain and generalize well across many tasks^37,38^.

Moreover, our dataset is also compatible with multiple stimulus annotations provided by the existing *studyforrest* project^39–41^. Because the visual stimuli for the MEG and fMRI datasets can be precisely aligned via the technique demonstrated in Fig. 2, the various non-speech related visual stimulus annotations from the German-dubbed fMRI stimuli in the *studyforrest* project can be used to complement the MEG data to explore the spatiotemporal dynamics underlying cognitive processing of annotated features. Moreover, although the MEG and fMRI auditory stimuli cannot be strictly aligned at the phoneme level due to the language differences, annotations describing semantic features (e.g. semantic conflict) for fMRI stimuli should also work for the MEG dataset.

Despite the importance of the aforementioned dataset as an extension of the *studyforrest* dataset in studying the spatiotemporal dynamics underlying cognitive processing in real-life contexts, the limitations should be acknowledged. First, the participants in our MEG data did not overlap with that in the *studyforrest* project. Therefore, the fMRI-MEG fusion can be only performed at the group level (i.e., across participants) instead of at the individual level (i.e., within participants), thereby hampering the ability to study the individual differences in the coupling between spatially localized networks and temporal dynamics. Second, the dubbed languages used in our dataset and the *studyforrest* project were radically different, limiting the application of the data in examining spatiotemporal dynamics of brain activity underlying auditory processing and language. In addition, time differences between the stimuli in the MEG and fMRI data should be treated with caution. Considering the lower sensitivity of fMRI signals to the exact timing than that of MEG signals, we recommend the use of MEG stimuli for fusing fMRI and MEG data.

## Acknowledgements

This study was funded by the National Key R&D Program of China (Grant No. 2019YFA0709503) and the National Natural Science Foundation of China (Grant No. 31771251).

## Code Availability

All custom codes for data preprocessing and technical validation are available at https://github.com/BNUCNL/MEG_Gump. Preprocessing was performed using MNE-BIDS v0.8 (https://mne.tools/stable/index.html), MNE v0.22 (https://mne.tools/stable/install/mne_python.html), fMRIPrep v20.2.1 (https://fmriprep.org/en/stable/), pydeface v2.0.0 (https://github.com/poldracklab/pydeface), and dcm2niix v1.0.20180622 (https://github.com/rordenlab/dcm2niix).

## Author contributions

X.L. conceived and performed the study and wrote the manuscript. Y.D. performed the study. H.X. contributed to the data collection. Z.Z. conceived and supervised the study and wrote the manuscript.

---

**Supplementary Figure 1.**
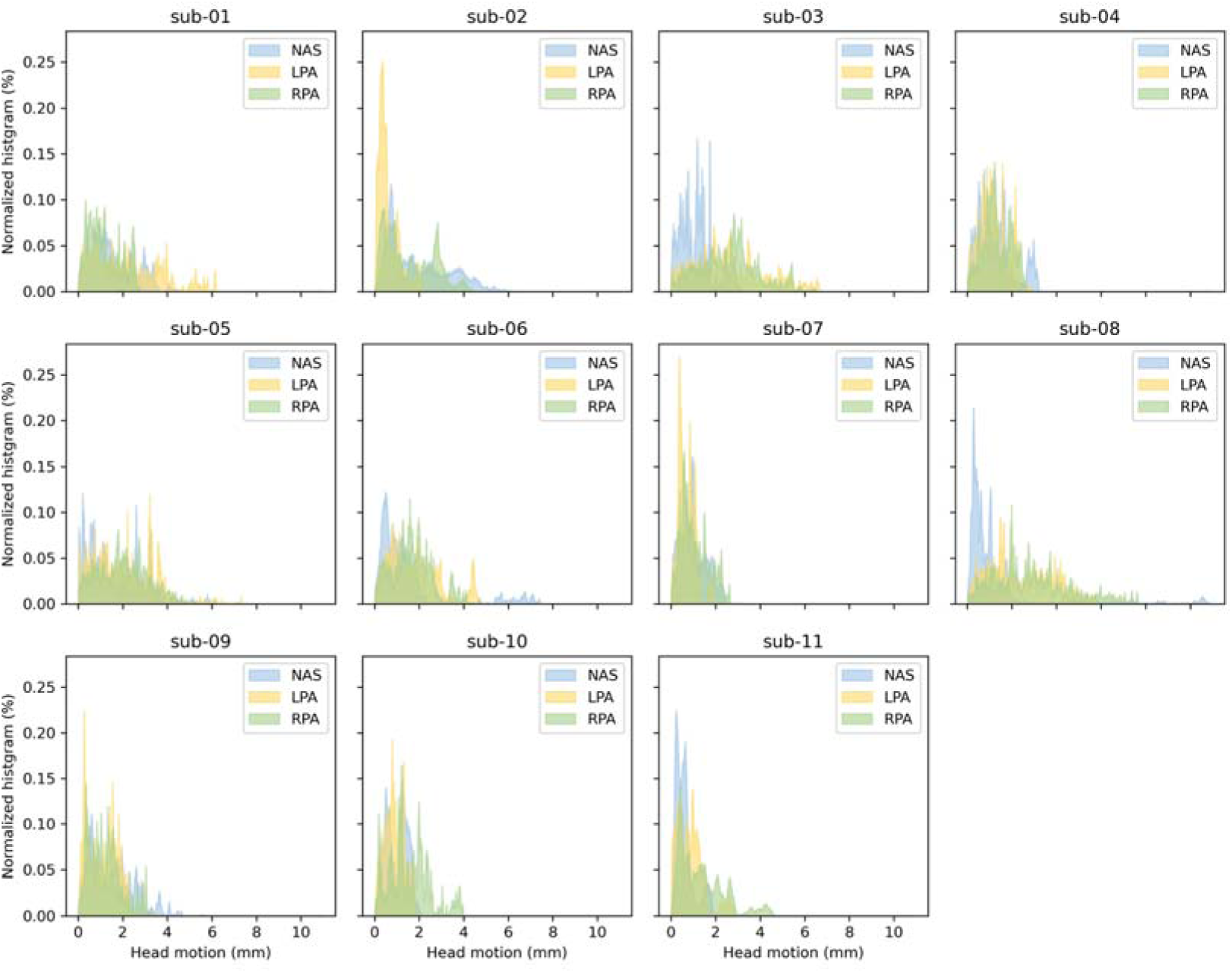
Head motion magnitude from each individual participant. The density histogram of motion magnitude calculated for three fiducials (NAS, LPA, RPA) were plotted for all samples, across all runs for each participant.

